# Integrative epigenomic analysis uncovers asymmetry of enhancer activity in *Brassica napus*

**DOI:** 10.1101/2025.10.31.685802

**Authors:** Silvia F. Zanini, Kevin Rockenbach, An Nguyen, Kübra Arslan, Gözde Yildiz, Rod J. Snowdon, Agnieszka A. Golicz

**Affiliations:** Department of Agrobioinformatics, IFZ Research Center for Biosystems, Land Use and Nutrition, Justus Liebig University, Heinrich Buff Ring 26-32, 35392 Giessen, Germany; Department of Plant Breeding, IFZ Research Center for Biosystems, Land Use and Nutrition, Justus Liebig University, Heinrich Buff Ring 26-32, 35392 Giessen, Germany

## Abstract

Non-coding regulatory regions are essential to the determination of gene expression and plant phenotypes. In this work, we investigated the cis-regulatory landscape of a winter type rapeseed, Express617, across multiple sample types. Combining chromatin accessibility, DNA methylation and gene expression, we annotated thousands of novel regulatory elements in the *Brassica napus* genome. Among those regions, we discovered and functionally characterized super-enhancers, observing an asymmetrical distribution of these regulatory elements favoring the Cn subgenome. Super-enhancer (SE) associated genes were found enriched in tissue identity and responses to stimuli related processes. We further establish and apply an in-silico validation pipeline for super-enhancers, integrating population-level expression analysis and machine learning (ML) models predicting gene expression levels. Almost 50% of the newly identified SE-associated genes had an observed expression higher than the expression levels predicted by the ML model. Moreover, structural variants disrupting super-enhancer elements correlate with a reduction of expression in the associated genes, both consistent with the positive effect of these regulatory regions. These results greatly expand the functional annotation of rapeseed and contribute to a better understanding of the link between regulatory elements and their target genes and processes, providing novel insights and targets for *B. napus* (epi)-genome editing strategies.

## Introduction

Plant development and its ability to adapt to external stimuli are determined by multilayered gene expression regulation, achieved by synchronization of chromatin organization and DNA accessibility. Main actors determining gene expression are transcription factors (TFs), which bind to cis-regulatory elements (CREs) based on the presence of a specific sequence pattern or motif and recruit additional co-factors to either promote or repress transcription of nearby genes. The ability of TFs to bind to their respective TFBS (transcription factor binding sites) is based on the DNA being accessible. DNA packaging into nucleosomes and/or high levels of cytosine methylation act as physical barriers to TF binding, rendering the regulatory region inaccessible and thusly inactive. These two features differ in their activity and stability, with DNA methylation being highly conserved throughout the life of a plant, with most methylation changes being due to the switch from vegetative to reproductive growth, but also as a consequence of stress exposure [1–5]. On the other hand, chromatin accessibility (or nucleosome-free chromatin stretches) has been shown as highly variable depending on the specific cell type, cell/life stage and presence of stimuli [6–11]. In addition to low DNA methylation and chromatin accessibility, histone modifications can further substantiate the regulatory role of the identified regions and help discern between CRE categories [12].

However, identification of non-promoter regulatory regions from their DNA sequence alone often results in misleading results and false positives, thus, including as many epigenetic features as possible (e.g. chromatin accessibility, DNA methylation, histone modifications, chromatin conformation) is, although both resources and experimentally demanding, paramount [13]. These regulatory regions are referred to as enhancers or silencers based on their positive or negative effect on associated (or cognate) genes expression, respectively, and can be found both distally or within introns of their target genes. Although silencers remain largely uncharacterized in plants, several examples exist proving the contribution of enhancers to interspecific variation and proposing them as key drivers of domestication in crop species [14–20]. Enhancers have been linked to the fine-tuning and spatio-temporal expression of several genes, ranging from developmental functions to specific stimuli responses. In bread wheat, *pathogenesis related 10 (PR10)* homeologs were found differentially expressed across the three subgenomes based on the differential expression of associated enhancers, with the A subgenome PR10 being targeted by a transcribed enhancer and exhibiting higher expression compared to B and D’s, associated with untranscribed enhancers [21]. In maize, several enhancers were found involved in modulating genes involved in leaf development and were hypothesised to be the result of TIR transposable elements domestication into distal regulatory sequences [22]. In potato, a 200bp regulatory element located in the second intron of a vacuolar invertase gene was deleted to prove its function as a transcriptional enhancer and its role in cold-induced sugar accumulation in post-harvest potatoes [23]

Recently, Zhao and colleagues have identified and validated the first examples of super-enhancer (SE) elements in plants [24]. These elements are known in mammals as key regulators of cell identity, both for healthy and diseased cell [25]. Consistently, SEs were shown to play a major regulatory function in organ development and tissue identity in Arabidopsis, with a partial disruption of the SE sequence leading to reduced expression levels of associated genes and distinct phenotypic changes [24].

*Brassica napus* is an allotetraploid derived from the hybridization of *B. rapa* (A subgenome) and *B. oleracea* (C subgenome), resulting in a 2n=38 (AACC) ∼1Gb genome. Given the relevance of *B. napus* as a major oil crop, especially in Europe, we aimed to shed light on the function of these non-coding regions and whether they could be hiding regulatory roles. To that end we chose Express 617, a winter oilseed rape (WOR) accession extensively used for rapeseed breeding efforts in continental Europe [26]. To that end, we assessed chromatin accessibility and DNA methylation, both common markers of functional genomic space, coding and not, in five distinct plant sections/tissues, including: leaves, roots, seedlings, immature flower buds and immature siliques. These were selected to represent the most diverse tissue/cells/life stages building towards a comprehensive blueprint of the regulatory space in rapeseed. Altogether our results confirm the increased power of multi-omics approaches in regulatory regions/CREs identification and identify a subset of high-potential candidates for fine tuning traits. We identify and predict how super-enhancer-like elements regulate both developmental and stimuli responses in *B. napus* by increasing associated genes’ expression in a tissue-specific manner. We observe an asymmetric distribution of these regions across subgenomes, with an enrichment of SEs in Cn compared to An. Finally, we provide *in silico* validation of their regulatory roles by integrating eQTL analysis and deep learning model predictions. Our multi-faceted approach to mapping out the regulatory landscape of a high-value crop plant paves the way towards applications of epigenetic multi-omics studies in trait discovery and refining.

## Results

### Open chromatin landscape of *Brassica napus* genome

To investigate the active chromatin regions and cis-regulatory elements in WOR we selected five diverse tissues/organs of *B. napus* Express617: seedlings, leaves, roots, immature flower bud bundles (referred in the rest of the text and figures as flowers for brevity) and immature siliques (siliques). Two different approaches were used to generate ATACseq libraries: eight samples were generated from semi-crude isolated nuclei, encompassing four sample types (leaves, seedlings, flowers and siliques), averaging 23x coverage per library. Root semi-crude libraries failed to meet the minimum criteria during data QC and were consequently discarded, and new root tissues were sent for FACS-based nuclei isolation and ATACseq (50x coverage per library). An average of 31,000 open chromatin regions (OCRs) were identified per sample type, with siliques being the most accessible (46,500 OCRs corresponding to 21Mb) and seedlings being the least (16,400 OCRs corresponding to 6.4Mb) (**Figure 1c**). Although mostly ranging from 250 to 700 bp, the average length of OCRs varied significantly among tissues, with leaves (average 369 bp, median 303 bp), seedlings (average 392 bp, median 314 bp), flowers (average 411 bp, median 339 bp), siliques (average 463 bp, median 378 bp) and roots (average 613 bp, median 477 bp) (**Figure 1d**). Concordant with their number and length differences, the total space occupied by OCRs varied across tissue, from 0.72% of the genome accessible in seedlings, 0.98% in leaves, 1.49% in roots, 2.22% in flowers, up to 2.42% in siliques.

**Figure 1.**
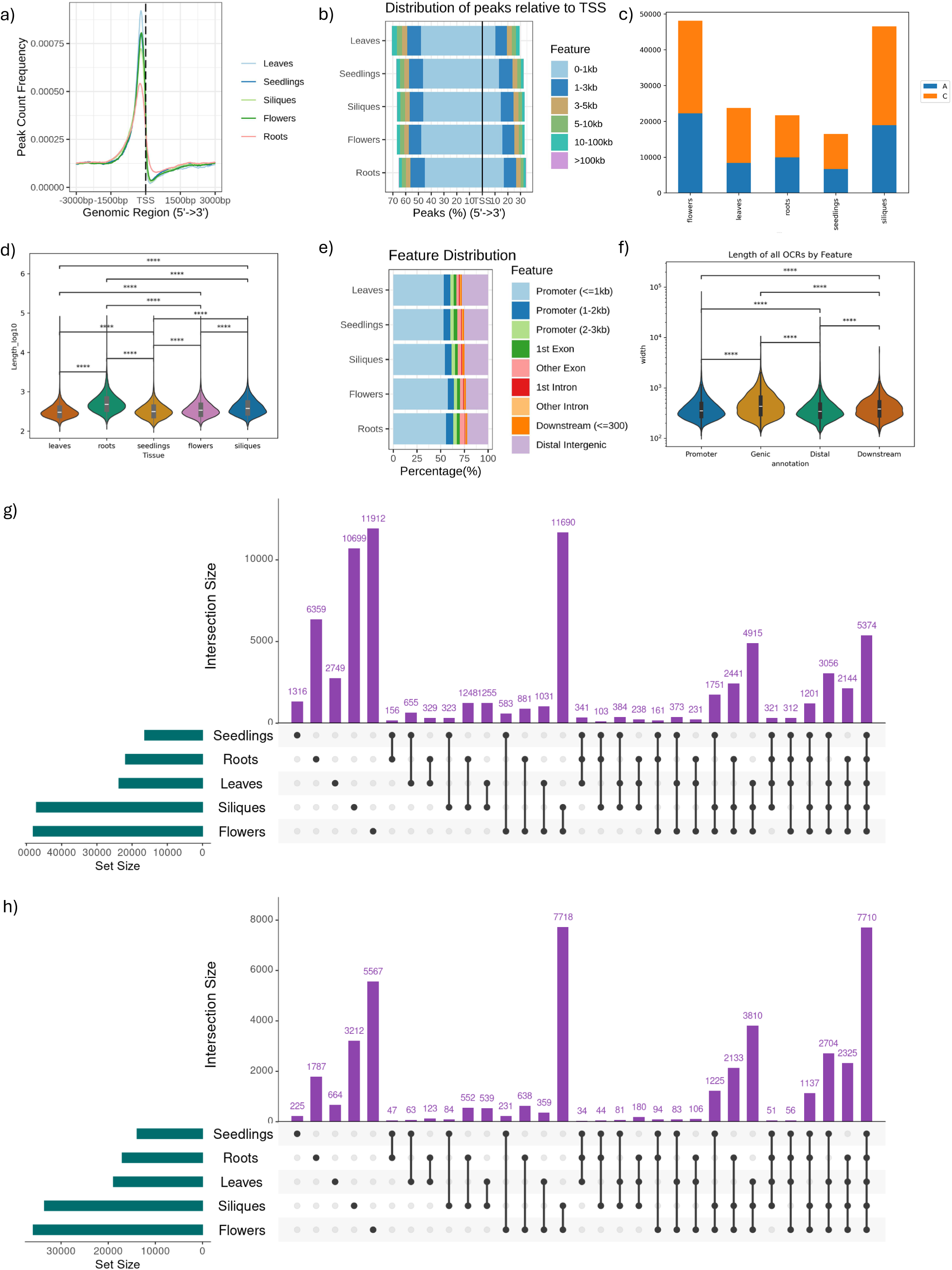
*Brassica napus* open chromatin regions. a) Frequency of ATACseq peaks at genes transcription start site (TSS), b) distance of ATACseq peaks to TSS, c) number and subgenome distribution of open chromatin regions (OCRs) . d) length of OCRs by tissue, e) Annotation of features of ATACseq peaks f) length of OCRs by feature annotation, g) conservation of ATACseq peaks across tissues, by reciprocal 50% overlap, h) conservation of ATACseq peaks across tissues, by conservation of nearest (cognate) genes. Significance test: Mann-Whitney U test.

The distribution of ATACseq peaks was enriched in the transcription start site (TSS) region for all investigated tissues, with approximately 60% of peaks being found 1kb or less from the TSS (**Figure 1a-b**). In concordance with the highly dynamic state and tissue-specificity of open chromatin, only 5,374 OCRs were consistently identified in all tissues based on region overlap (reciprocal minimum overlap of 50%, **Figure 1g**). Interestingly, a higher degree of conservation was observed when comparing chromatin accessibility profiles based on closest (cognate) gene conservation, resulting in 7,710 genes being near OCRs in all tissues assessed (**Figure 1h**).

All OCRs were annotated in relation to the corresponding genomic space and divided into “genic” (overlapping > 1bp with gene body), “promoter” (overlapping with the region from 1bp to 3kb upstream of TSS) or “intergenic” (not overlapping annotated features) regions (**Figure 1e**). On average, 65 % of OCRs were annotated as “promoter” in each tissue, with an additional 10% “genic” and the remaining 25% “intergenic”. OCR size was found to vary depending on its genomic location/annotation, with distal or promoter OCRs being on average shorter than genic and downstream regions (**Figure 1f**).

Transcription factor binding motifs (TFBM) were identified and annotated *in silico* in all OCRs with MEME FIMO using the curate core Jaspar 2024 plants motif database. *In silico* prediction of TFBM is intrinsically prone to high false positive rates due to the limited specificity of TF affinities and especially in the case of input sequences longer than 500bp. Although relying on stringent q-values prevents several motifs from being annotated entirely, we compare annotation rates between OCRs and randomized regions of the same length and number, observing a strong increase (5x) in motifs annotated in OCRs, on average 4.6 motifs per 1kb, compared to randomized regions, averaging 0.9 motifs/kb. A less stringent prediction (*P=*0.0001) resulted in an unrealistic number of TFBS annotated (>20motifs/100bp) but still exhibited a similar trend when comparing OCRs and randomized regions (1.6x increase).

### DNA methylation profiling supports presence of proximal and distal regulatory regions

Whole genome bisulfite sequencing was carried out for the three most representative tissues (leaves, roots and siliques), generating approximately 33 Gb of data per replicate, with 2 biological replicates per sample type.

Genome wide methylation rate was approximately 44-46% in CG context, 3-4% in CHH and 16-20% CHG, consistently across tissues. Gene-based methylation status assessment confirmed the expected decrease in methylation rate in gene bodies compared to non-coding regions in all tissues and contexts, especially CG (**Figure 2a, Supplementary Figure 1**). Similarly, methylation rate of OCRs was calculated in each of the three contexts and confirmed to negatively correlate with chromatin accessibility (**Figure 2b, Supplementary Figure 1**). To further corroborate the distal OCR identification as CREs, we assessed the methylation status of OCRs > 3kb from gene bodies and observed an appreciable decrease in methylation levels in these regions (**Figure 2c, Supplementary Figure 1**).

**Figure 2.**
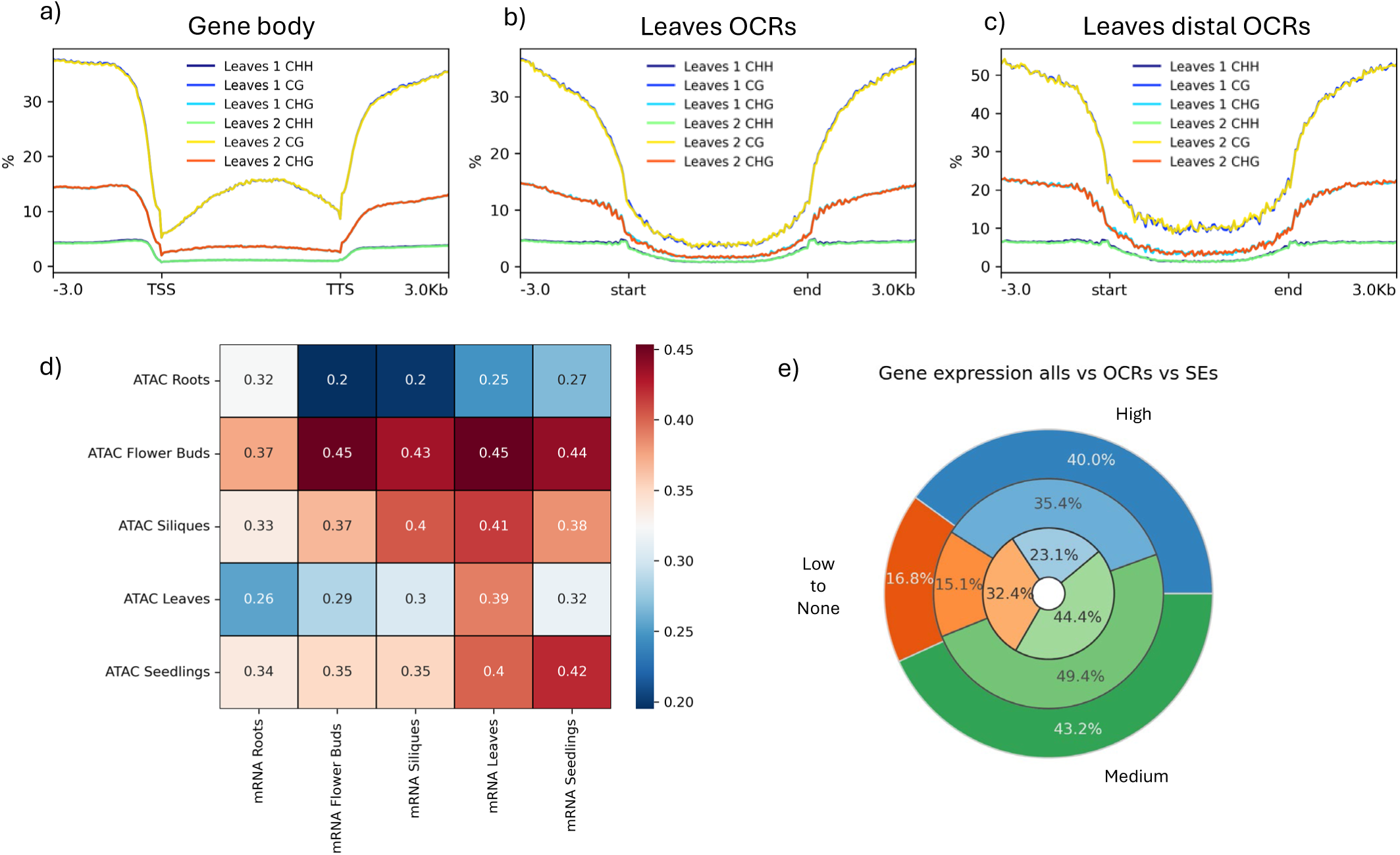
Methylation rate and gene expression assessment across open chromatin regions. a-c) DNA methylation in all 3 contexts (CHH,CHG and CG) across genic regions (a), OCRs (b) and distal OCRs (c) for leaves samples. d) Spearman correlation coefficient matrix based on read coverage of ATACseq libraries and mRNAseq libraries calculated across extended promoters (3kb) and genic regions. e) Gene distribution based on expression (TPM) genome-wide( inner circle), OCRs associated (middle circle) and SE associated genes (outer circle). Genes were divided as following: “low to none” = 0 to 1 TPM , “medium” 1 to 10 TPM and finally “High” above 10TPM.

### Integration of gene expression confirms regulatory activity of OCRs

Finally, the relationship between chromatin accessibility and gene expression was assessed based on both raw and processed signals (**Figure 2d**). On average, each tissue matching achieved 0.4 correlation coefficient between data types (ATAC vs RNA), with flowers exhibiting the highest (0.45) and roots the lowest (0.32) correlations. The strongest correlation within data types was observed for the two reproductive tissues, flowers and siliques (0.89 ATAC and 0.87 RNA), while roots were the most distinct tissues, with 0.83 RNA and 0.72 ATAC correlation coefficient when compared to the closest tissue, seedlings (**Supplementary Figure 2**).

Furthermore, for each tissue genes were divided into none-low (<1TPM), medium (1-10TPM) and high expression (> 10 TPM). Approximately 32.4% of genes were assigned to the lowest expression tier, 44.4% to the medium and 23.1% to the highest across tissues (**Figure 2e**). Genes were then subset if neighboring OCRs, with near OCR-genes found enriched in the highest (35.4%) and mid (49.4%) expression tier and depleted in the lowest (15.1%), consistent with a positive regulatory effect of the identified OCRs.

### *Brassica napus* genome encodes hundreds of super-enhancer elements

Among all accessible chromatin, we focused on super-enhancer (SE) elements, defined according to the criteria utilized by Zhao and colleagues as top 2.5% non-core promoter (500-0bp from TSS) OCRs by length, and known to have a strong positive effect on neighboring genes expression in Arabidopsis [24]. Here, we identified 381 candidate super-enhancers (SEs) in leaves, 205 in roots, 649 in siliques, 290 in seedlings and 668 in flowers (**Figure 3a-d**). Genes were subset if neighboring candidate SEs and classified once more based on their expression (low to none, mid or high), denoting a further increase in the ratio of both high and low expressed genes and a decrease in medium expressed ones when compared to genes neighboring OCRs, in concordance with the expected stronger regulatory effect of these SEs (**Figure 2e**).

**Figure 3.**
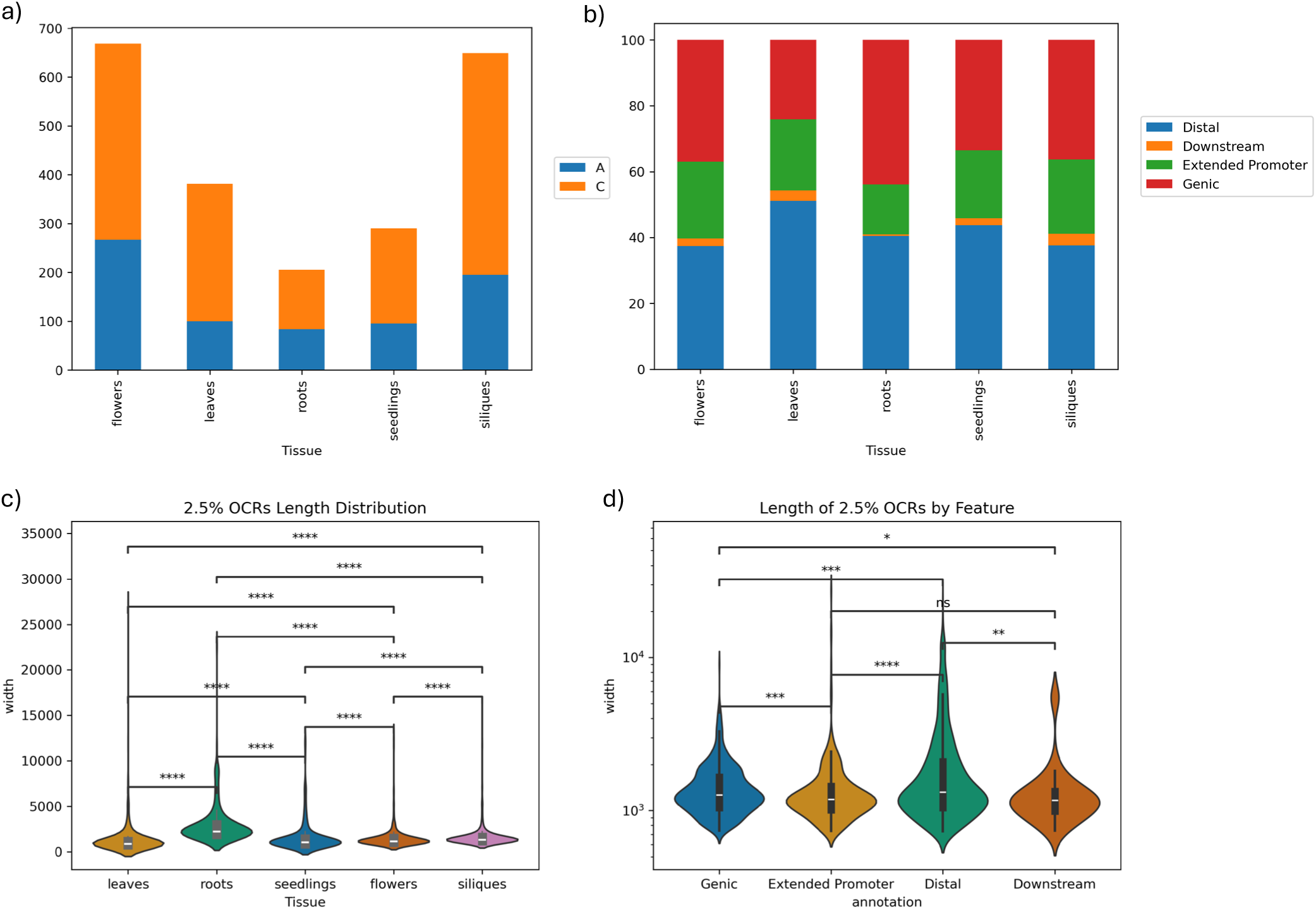
Super-enhancers characterization in *Brassica napus*. a) number and subgenome distribution of candidate SEs across the five tissues. b) annotation distribution of candidate SEs across the five tissues, c) length distribution of candidate SEs across the five tissues. d) length distribution of candidate SEs based on their annotation. Significance test: Mann-Whitney U test.

While the proportion of SEs in the extended promoters (3kb from TSS) of their cognate genes is relatively stable across tissues (∼20%), leaves show a higher proportion of upstream distal SEs (∼50%) compared to the other tissues (∼40%, **Figure 3b**). Additional tissue specific variability was observed in their length distribution (longest in roots, **Figure 3c**) and in the average distance of SEs up to 10kb downstream of their cognate genes in leaves (**Figure 4c-d**). Similar patterns were observed in the distribution profiles of SEs distances both up and downstream their cognate gene, within most SEs being found within 2kb of their associated gene (**Figure 4a-b**).

**Figure 4.**
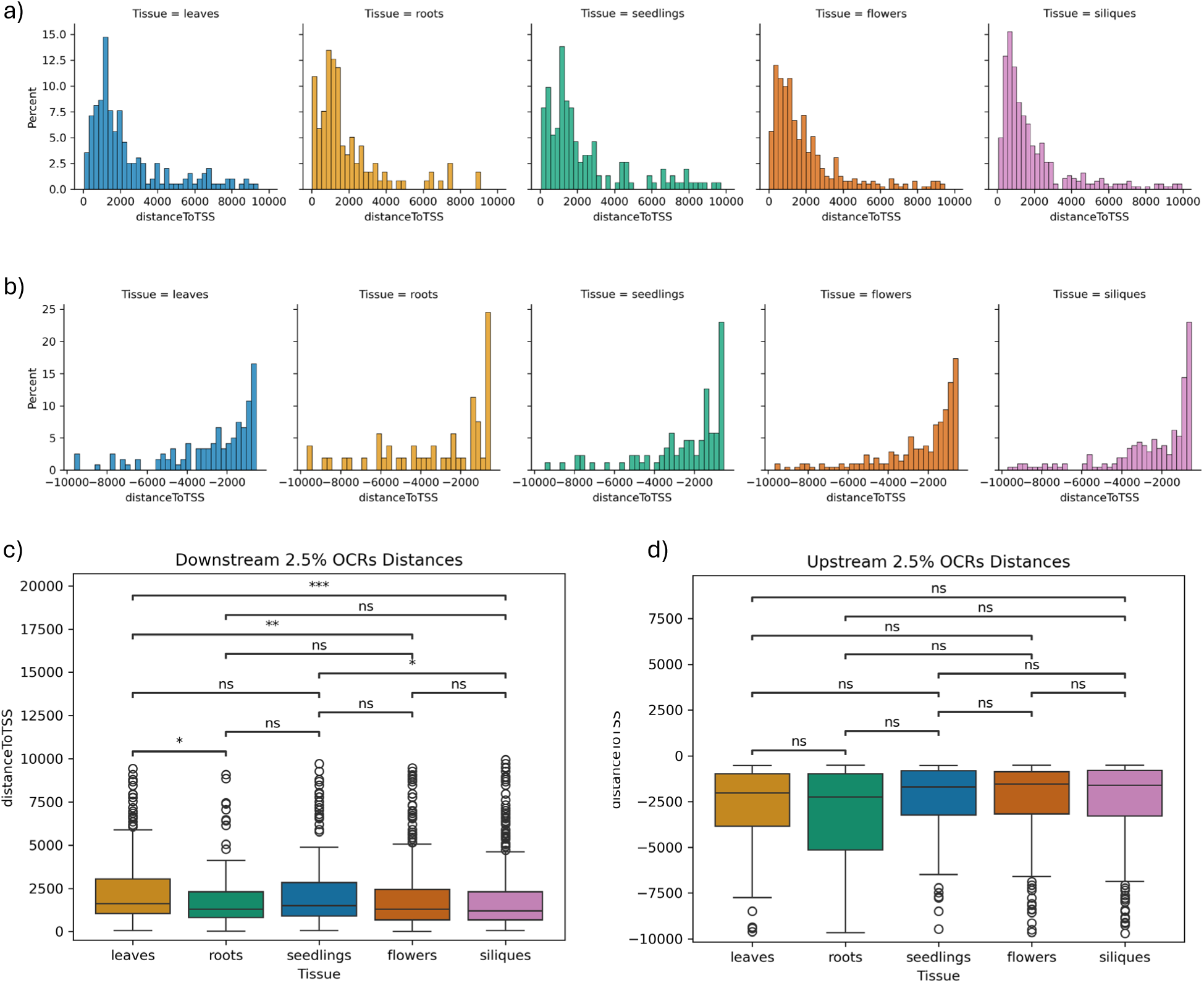
Super-enhancers length and distribution across tissues. a) downstream super-enhancers (SE)s distance to gene transcription start sites (TSS) distribution (subset of SEs up to 10kb downstream, which also include genic SEs) b) upstream SEs distance to gene TSS (subset of SEs 500 bp up to 10kb upstream) , c) downstream SEs distance to gene TSS. d) upstream SEs distance to gene TSS (subset of SEs 500 bp up to 10kb upstream) . Significance test: Mann-Whitney U test.

To understand the effect of active super-enhancers on their cognate genes, we compared expression levels of their cognate genes in the corresponding tissue with expression levels in the other four samples. We observed higher expression in the SE-associated subset compared to non-SE in the corresponding tissue, but no or reduced effect in the other tissues, further indicating the (tissue-specific) regulatory role played by these regions (**Figure 5b-f**). To assess whether the tissue specificity observed in SEs is also mirrored in their associated genes, we investigated gene tissue specificity metrics (TAU) across all annotated *B. napus* genes. TAU values were then subset based on whether the corresponding gene has an open core promoter (0 to 500bp upstream of TSS) but no SE assigned, or both open core promoter and an assigned SE. No significant difference in tissue specificity values was observed between the two gene subsets (**Figure 5a**).

**Figure 5.**
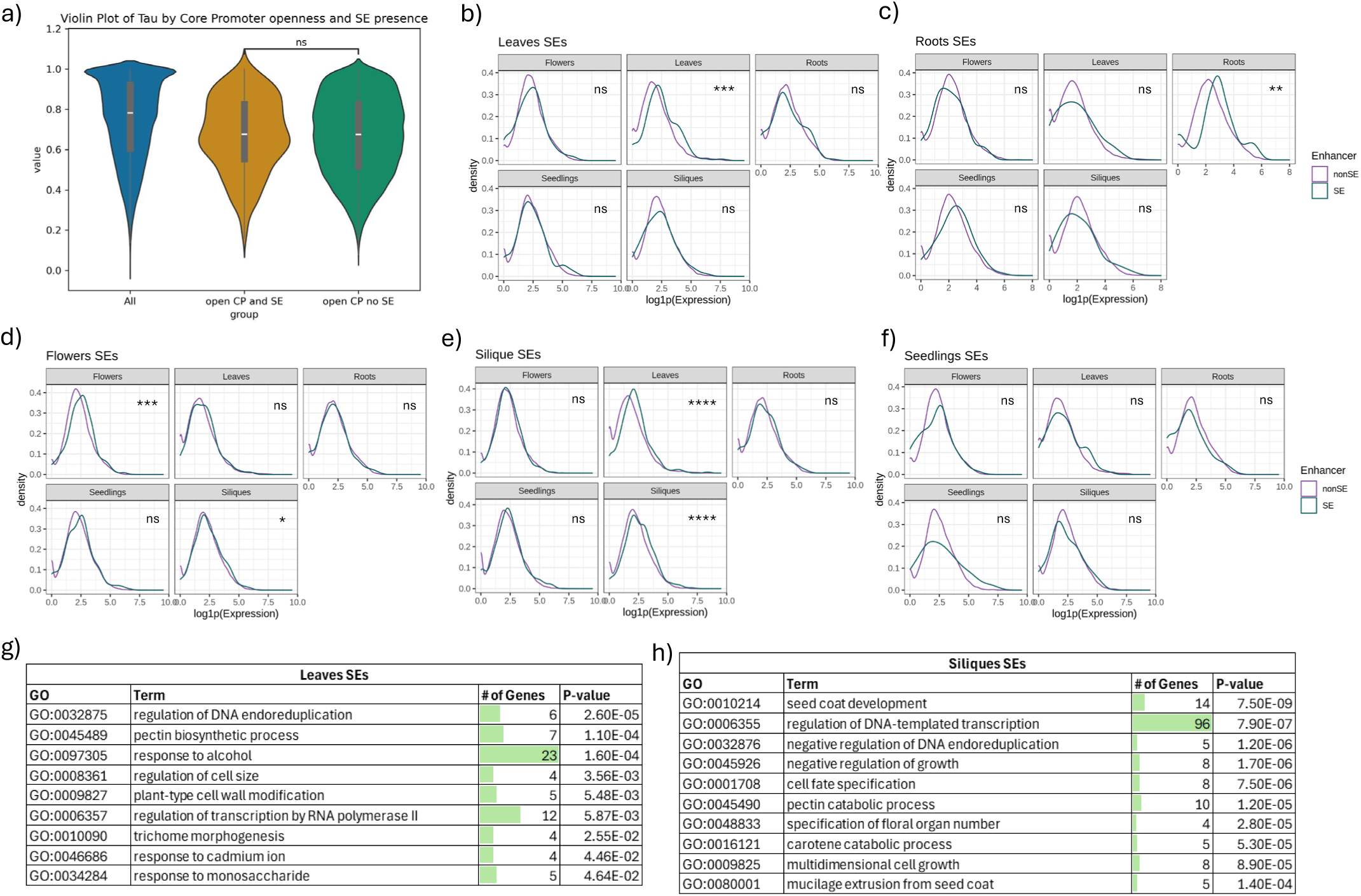
Super-enhancer associated genes. a) Gene tissue specificity (TAU) values based on their expression across all tissues, subset based on their core promoter (CP) being accessible and association (or not) with super-enhancers (SEs) . Significance test: Mann-Whitney U test. b-f) effect of SEs on cognate gene expression in the corresponding tissue compared to the other four. Significance: Mann-Whitney 1 sided test , g-h) top Gene Ontology Enrichment of SE-cognate genes in leaves (g) and siliques (h).

### Functional analysis of SE-cognate genes points to enrichment of developmental processes

Gene Ontology Enrichment (GOE) analysis highlighted an enrichment in leaves SE-cognate genes versus all OCRs cognate genes of terms related to leaf specific processes such as regulation of DNA endoreduplication and trichome morphogenesis, as well as response to stimuli including alcohol, cadmium and monosaccharide (**Figure 5g**). Consistent with the early growth stage of these samples, seedlings SE-cognate genes were enriched both with terms related to plant growth and organ development, tropism and response to stimuli (**Supplementary Figure 3**). Conversely, SE-cognate genes from the two reproductive stage samples exhibited enrichment in terms related to seed coat development and mucilage extrusion, and cell fate specification and growth (**Figure 5h, Supplementary Figure 3**), indicating a key role of SEs in coordinating responses to a wide range of stimuli and tissue determination, especially in the reproductive stage of *B. napus*. Finally, to investigate the role of SE associated genes we performed weighted gene co-expression network analysis (WGCNA) across 36 RNAseq samples from [27] and our newly generated 5 RNAseq samples/tissues. This analysis resulted in 35 modules, out of which 29 included SE associated genes. We did not observe a significant variation in gene connectivity network-wide (**Supplementary Figure 4**) when comparing SE or OCR associated genes to the remaining closed-chromatin genes. When comparing only within modules we observed diverging trends with some modules exhibiting a reduction in gene connectivity for open chromatin (OCRs and/or SEs) associated genes and vice versa.

### Super-enhancer elements are asymmetrically distributed across the subgenomes

We assessed the expression levels of all genes in the An and Cn subgenomes, observing higher levels of genes expression from the An subgenome across all five tissues and a slight increase in tissue specificity in Cn genes (**Figure 6d-e**). We further investigated this subgenome asymmetry in the epigenetic datasets, confirming an overall increase in methylation levels in the C subgenome both in gene bodies and regulatory regions (**Figure 6b-c**), also reported in previous studies [28–30]. Conversely, more open chromatin regions and candidate super-enhancers were found in the C subgenome (**Figures 1c, 3a**), which is consistent with the presence of more protein coding genes annotated on these chromosomes (approximately 47.6% genes annotated are in the An subgenome versus 52.4% in Cn). We identified approximately 42.3% OCRs in An versus 57.7% Cn (**Figure 1c**), and 33.8% SEs in An compared to 66.2% Cn (**Figure 3a**), denoting an increase in asymmetry among the longest accessible chromatin regions [31]. To elucidate whether this bias for the C subgenome in chromatin accessibility is due to uniquely Cn genes or an imbalance in homeologs, we identified homeolog pairs based on collinearity between subgenomes (**Figure 6a**). We identify 11,800 genes unique to the An subgenome and 16,100 to Cn. Of these, 4,300 unique An genes were associated to OCRs compared to 6,200 unique Cn (χ^2^= 6.61, *P=* 0.01). This enrichment of accessible Cn unique genes was also observed in the super-enhancers associated genes subset (227 unique Cn vs 119 unique An genes), both when compared to genome-wide (χ^2^= 8.89, *P=* 0.0029) and to the OCR-associated subset (χ^2^=6.37, *P=* 0.0116) (**Figure 6g**). GOE analysis highlighted different roles for these SE-associated subgenome-specific genes, with terms related to RNA processing, cell size/growth regulation and catabolic processes in SE-An unique genes, whereas terms related to photosynthesis, seed coat and response to stimuli were found enriched in SE-Cn unique genes (**Figure 6f**).

**Figure 6.**
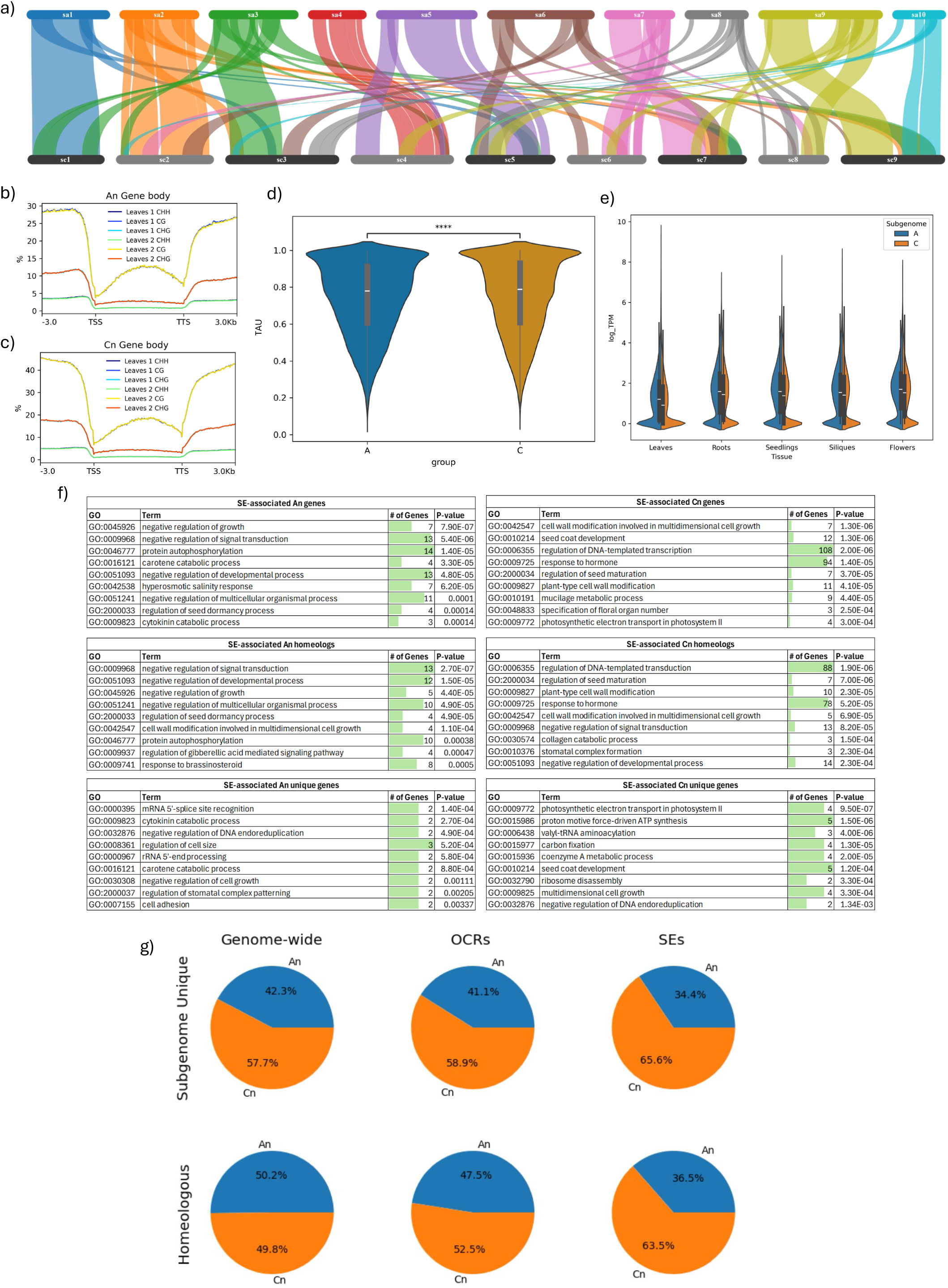
Subgenome asymmetry in *Brassica napus*. a) collinearity between A and C subgenomes. Blocks with a final score of minimum 9047 were visualized with SynVisio , b-c) DNA methylation across An (b) and Cn (c) gene bodies, d) tissue specificity score (TAU) of An and Cn genes, Significance test: Mann-Whitney U test, e) gene expression distribution (log transformed) across tissues and subgenomes, f) GO terms enriched in subgenome super-enhancer (SE)-associated genes, g) percentages of subgenome-unique or homeologous genes: genome-wide, OCR-associated and SE-associated.

### Integration of SEs with machine learning-based gene expression prediction validates their functionality

To further assess the predicted SE function and effects, we combined the newly generated data with a machine learning (ML) based approach utilized to predict gene expression from only DNA sequence [32] (**Figure 7a**). When investigating the correlation between predicted and observed expression levels, we observed a population of genes whose expression was under-predicted by the model. We observed in Rockenbach et al. [32] the enrichment of super-enhancer associated genes among poorly predicted ML expression. The *n*emo ML model was trained based on extended (6.2 kb) promoter and terminator regions but assigned the highest importance (by a large margin) to sequences ±1kb around the TSS or TTS, which mirrors the conserved and integral role of promoter and terminator regions and their abundance in the genome. Although training regions encompass a substantial proportion of our candidate SEs, we expect the model to not have enough SE training examples compared to promoters/terminators to fully capture and account for their effects. Genes that were found either over or under predicted by the model when compared to actual gene expression from our RNA-seq datasets were further investigated as putatively distally regulated. A total of 54,536 genes were matched between the two references versions and had predicted expression by the model. Expression levels were then compared between predicted and observed TPM and finally genes were divided based on the concordance between these values, resulting in 27,365 (50%) genes which had expression levels correctly predicted compared to the observed, 15,164 (28%) were underpredicted and 12,007 (22%) overpredicted by the model compared to the real/observed expression (**Figure 7c**).

**Figure 7.**
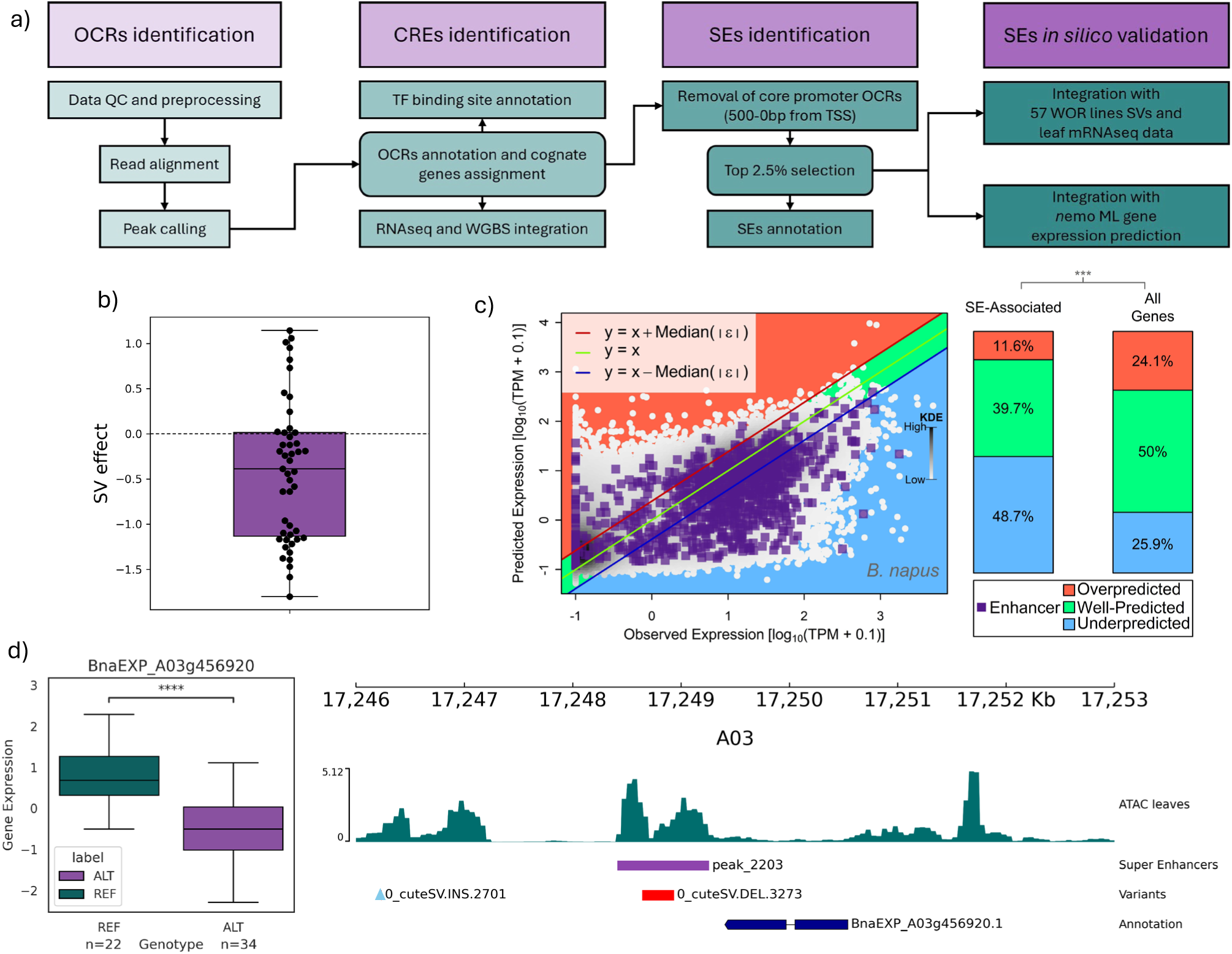
*in silico* validation of rapeseed super-ehnacers. a) overview of in silico approaches for the identification and validation of super-enhancers (SEs). b) predicted effect of structural variants disrupting SEs on the associated genes’ expression. c) machine learning model prediction accuracy across all genes and all SE-associated genes. d) example of gene expression variation in the presence of a deletion disrupting the neighboring SE. Significance test: Mann-Whitney U test.

We identified a prevalence of underpredicted genes in the SE cognate gene sets, with 369 (39.7%) correctly predicted SE-cognate genes, 453 (48.7%) underpredicted and 108 (11.6%) overpredicted. This observation is consistent with our initial hypothesis of a positive regulatory role of these candidate SEs, which is not accounted for in the ML model and as such results in an underprediction of the associated genes expression. Poorly predicted SE-cognate genes included transcription factors of the AGL (AGAMOUS-Like) family, involved in floral and fruit development as well as several genes involved in disease resistance and stress tolerance, including resistance to *Botrytis cinerea*, mildew, cold, drought, salt and heat.

### Structural variants located in SE elements putatively impact associated gene expression

To further validate the positive effect of SEs on gene expression, we assessed how naturally occurring sequence variation in these regions is reflected on the associated gene expression (**Figure 7a**). To that end, structural variant detection and genotyping were performed as previously described in Yildiz et al [33], resulting in the identification of a total of 77,142 SVs, including 37,348 insertions and 39,667 deletions across 57 rapeseed elite inbred lines. We then investigated the presence of these variants in our OCRs and specifically in the set of leaves-SEs, to match the plant tissue sampled for RNAseq. We observed an enrichment of SVs in OCRs (**Supplementary Figure 5**) and identified 62 variants overlapping a total of 57 SEs. Gene expression across the 57 samples was calculated for all cognate genes and related back to genotyping information to assess whether the presence of structural variants in the associated SE matches a reduction in gene expression, which would corroborate the positive regulatory effect of these SEs. Although the sample size was too small for a well resolved association study (e.g. eQTL), we still observed statistically significant variation in gene expression in the presence of SE-SVs in 19 genes (test Mann-Whitney two sided), out of which 16 led to significantly reduced expression in the presence of the SVs. An additional 17 out of 27 associations did not reach statistical significance but were still showing a trend consistent with a positive regulatory role of the SEs and a deleterious effect of the SV disruption (**Figure 7b**).

Although this *in silico* validation approach only included a modest set size, we successfully identified four SE-genes pairs with a significant downregulation in gene expression in the presence of -SVs disrupting the associated super-enhancer. These included a probable aquaporin linked to drought tolerance in Arabidopsis [34], two brassinosteroid-related proteins and finally BnaEXP_A03g456920, a BOI-related gene (BRG) involved in resistance to *Botrytis cinerea* (**Figure 7d**).

## Discussion

As a major oil crop, previous research has predominantly focused on the role of epigenetic mechanisms in rapeseed resulting in changes in seed oil content [35–37] and its asymmetric subgenomic chromatin accessibility and architecture [31]. However, a more comprehensive assessment of regulatory mechanisms encompassing both vegetative and reproductive stages of plant growth was yet to be achieved for one of the major European winter type references, Express 617. Additionally, rapeseed has a gene dense genome, with approximately 100 thousand transcripts and contains 37% repetitive regions. Majority of natural genetic variation occurring in *B. napus* is found in non-protein coding regions, as SVs/SNPs within coding sequences tend to have a starker impact on gene function and are subject to stronger selection pressure. Nonetheless, GWAS and eQTL studies in several models and crops species, including *B. napus* itself, have shown that these non-coding variants can also impact gene expression and/or modulate agronomically relevant traits [33, 35, 38–42]. In addition, these variants tend to lead to milder effects compared to their coding counterparts, providing the ability to fine-tune traits.

In the present work we investigated both conserved and tissue-specific epigenetics and transcriptomics profiles in five diverse tissues, generating a comprehensive map of open/accessible chromatin spanning the entire life cycle of the plant. We observed a modest proportion of conserved open chromatin regions (OCRs) across all samples, with most regions being only detected in one out of five samples except for the reproductive tissues analyzed, with more than half of OCRs being shared among flower buds and siliques. We also noted how genes associated (or closest to) OCRs were appreciably more conserved across samples, suggesting that different regulatory regions modulate the same gene in a tissue-dependent manner.

DNA methylation has been a marker for functional regions of genomes regardless of their active state, as it is overall stable across tissues and life stages and it associates with both coding and regulatory (CREs) regions. Approximately a third of rapeseed genome is unmethylated, so relying on WGBS data alone would not provide the resolution necessary to elucidate complex epigenetic processes. Nevertheless, low to no DNA methylation is a requirement for most transcription factors to bind to the DNA and perform their regulatory functions. As such, we assessed DNA methylation rate across three samples and observed a noticeable decrease in methylation in all three contexts (especially CG) in open chromatin regions, further substantiating their classification as CREs.

Finally, we corroborate the predicted regulatory role and effect of these newly annotated CREs by integrating gene expression data and observed that genes associated with OCRs were found to be on average higher expressed across tissues. However, the accessibility of chromatin only moderately correlates with gene expression (Pearson correlation coefficient ∼0.4), indicating that there are additional processes and layers of regulation beyond the ones assessed in this work.

We then focused on the longest OCRs not overlapping core promoters (500-0bps from TSS), as these were previously categorized as “super-enhancers” (SEs) in *A. thaliana* and were shown to be associated with genes involved in tissue identity and development, mirroring the role of SEs in mammals [24]. In *B. napus*, our analysis revealed an enrichment of SE-cognate genes related to seed development and floral organ number and identity in the two reproductive tissues analyzed, while the three vegetative tissues exhibited both enrichment in tissue identity related genes, but also of ones related to responses to stimuli, ranging from nutrients to abiotic and biotic stress responses. Genes associated with SEs were found to be on average higher expressed than OCRs associated but did not show a significant increase in tissue specificity, suggesting that these bigger regulatory regions function as modulators rather than on-off switches.

We further assessed the enrichment or depletion of transposable elements in both OCRs and super-enhancers (**Supplementary Figure 6**), observing a significant depletion of TEs annotated in both these subsets and indicating that, while TE insertions are more likely to occur in accessible chromatin, their insertion disrupts chromatin organization of the region, leading to the immediate loss of active/accessible status. Consistently, we observe a significant overlap of SVs identified across 57 WOR genotypes with Express 617 open chromatin regions, further substantiating the notion that TEs, as main drivers of sequence variation, accumulate preferentially in accessible DNA [43].

Throughout this analysis, we observed an asymmetric distribution of expression and epigenetic data between the A and C subgenomes, with the A subgenome exhibiting a lower degree of DNA methylation compared to the C subgenome and on average higher gene expression. Although this is consistent with previous studies in rapeseed [28, 29, 44–46], we also observed a contrasting asymmetry regarding chromatin accessibility, with more OCRs being identified in the Cn subgenome compared to the An. A recent article by Li and colleagues [31] corroborates our findings, observing higher accessibility of Cn-unique genes and proposes as an explanation of this imbalance the H3K27me3 modification. Although lacking histone modification datasets in our approach, we also confirmed an unexpected increase in Cn genes and specifically Cn-unique genes across open chromatin regions, even more so among candidate super-enhancers. Comparing SE-genes GOs across subgenomes, we observed distinct enrichments between An and Cn SE-genes, with mostly terms related to negative regulation enriched in the An subset. Interestingly, the enrichment of GO terms related to seed coat development and photosynthetic processes in Cn-SEs is driven by Cn unique genes rather than homeologs. Similarly, cytokinin and carotene catabolic processes related terms were found enriched in An-SEs and specifically An-unique SE genes.

Combining our regulatory blueprint with available SV and expression data from 57 German rapeseed elite breeding lines enabled a first validation of these sequences’ regulatory roles, by assessing the impact of SE sequence variation on the expression of the associated gene. Although 57 is too small of a set to be suitable for a comprehensive eQTL analysis, we still identified several SVs falling within or across SEs and correlating with altered expression of nearby genes.

Overall, our data highlights the importance of tissue-resolved multiomics, as a golden standard for epigenetic investigations. We discovered both tissue-specific and conserved super-enhancer elements in *B. napus* and observed an asymmetric distribution across the two subgenomes, with the Cn subgenome containing majority of the candidate SEs, in contrast with the overall lower expression levels of Cn genes. We further developed and implemented a pipeline to validate *in silico* these sequences, elucidating their regulatory code and effect on associated genes. Moreover, this work vastly extends the functional annotation of winter oilseed rape genome and provides the foundation for further regulatory elements profiling, both in the direction of (simultaneous) single cell ATAC-and RNA-seq (e.g. Epi Multiome ATAC + Gene Expression from 10x Genomics), which would enable a higher resolution and cell type-and stage-specific profiles, and the inclusion of further growth conditions, especially in the presence of external stimuli such as adverse climate conditions or pathogen pressure.

## Materials and methods

### Experimental design and plant growth conditions

All sequencing libraries (ATACseq, RNAseq, WGBS and MicroC) were derived from the same tissues, when possible, or from tissues from the same batch of plants. All experiments were replicated twice, and biological replicates were generated from separately grown experiments/batches, apart from long read PacBio sequencing for genome assembly, where only one individual was sequenced.

Plants were grown under long day conditions in growth chambers and vernalized for 10 weeks at 8/16 hours day/night cycles at a constant 5°C. Five sample types were collected in liquid nitrogen and stored at -80°C until further processing, thee representing vegetative and two reproductive growth stages: leaves from 3.5-week-old plants, 9-day-old roots, 3-day-old seedlings, immature flower buds and immature siliques.

### Data generation

#### ATACseq

ATACseq libraries were generated following two separated approaches. All five sample types were used for in-house ATACseq library preparation, while the roots were also subjected to FACS-based nuclei isolation and ATACseq by Azenta (Azenta US Inc.). For in-house library preparation, we followed a modified protocol derived from [6, 47, 48]. Briefly, approximately 0.3-0.4g of frozen tissue powder was resuspended in 3mL of NIB (15 mM Tris-HCl pH8, 20 mM NaCl, 80 mM KCl, 250 mM sucrose, 0.5 mM spermine, 5mM 2-ME, 0.3% TritonX-100 and protease inhibitor cocktail) and incubated on ice for 10 min with constant agitation. The solution was then filtered through pre-wetted miracloth and then through a 40 μm filter. Semi-pure nuclei were isolated by applying the lysate on top of a 1.7M sucrose solution (20 mM Tris-HCl Ph8.0, 2 mM MgCl2, 2 mM EDTA, 15 mM 2-ME, 1.7M sucrose, 0.2% TritonX-100) and centrifuged at 2,200 x g at 4°C for 20 minutes. Pelleted nuclei were washed twice by resuspending the pellet in iNIB (15 mM Tris-HCl pH8, 20 mM NaCl, 80 mM KCl, 250 mM sucrose) and centrifuging at 1000 x g at 4°C for 10 min. Isolated nuclei were stained with DAPI and counted to create aliquots of approximately 50,000 nuclei each for tagmentation. Each aliquot was resuspended in a tagmentation mix composed of 2.5 5 μl Tn5 transposomes in 47.5 μl tagmentation buffer and incubated at 37 °C for 30 min at 300 rpm. The tagmented fragments were cleaned with Zymo DNA Clean and Concentrator -5 kit and amplified with NEBNext® Ultra™ II Q5® Master Mix and ATACseq custom primers. The optimal PCR cycle number was determined by qPCR after the first 5 cycles, resulting in 10-12 total cycles per library. Finally, amplified libraries were purified with Zymo Select-a-size DNA Clean and Concentrator to remove all fragments below 100bp (primers) and sequenced.

#### RNAseq

RNA was extracted from all five sample types with Quick-RNA™ Plant Miniprep (Zymo Research) according to manufacturer instructions, including the optional on column DNase treatment. Quantity and purity of RNA was assessed with Qubit RNA BR Assay Kit and Nanodrop (ThermoFisher Scientific), RNA integrity with the RNA 6000 Nano Kit for 2100 Bioanalyzer Systems (Agilent). Samples were aliquoted and shipped on dry ice to Novogene Europe (Cambridge, UK) for library preparation and sequencing with strand specific Illumina 150PE.

#### WGBS

As the samples most representative of the full set, leaves, roots and siliques were subjected to whole genome bisulfite sequencing (WGBS) for DNA methylation assessment. Briefly, DNA was extracted with BioSprint 15 and BioSprint DNA Plant Kits (Qiagen), quality, quantity and fragment sizes were confirmed as described above before sending to Novogene Europe (Cambridge, UK) for library preparation and sequencing.

#### Long read PacBio HiFi sequencing

HMW-DNA was extracted from young *B. napus* Express 617 leaves with the Nucleobond HMW DNA extraction kit (Macherey-Nagel), following manufacturer instructions. DNA quality quantity and fragment size was evaluated as described above, before sending the sample to Novogene Europe (Cambridge, UK) for library preparation and sequencing on the Sequel PacBio instrument.

### Genome assembly

Long read DNA sequencing data was generated with PacBio HiFi and assembled with Hifiasm (v0.25) [49]. DNA fragments from organisms not belonging to Viridiplantae were initially classified and removed from the assembled contigs using a combination of tools, including Kraken2 v2.1.3 [50](core_nt database) and TaxonKit v0.20.0 [51]. To identify contigs originating from organellar DNA, the resulting contigs were then used as queries in BLASTn v2.14.1+ [52] against a database constructed from the chloroplast (NC_016734.1) and mitochondrial (NC_008285.1) DNA sequences of *Brassica napus*. Contigs with query coverage and identity greater than 90% to organellar DNA were excluded. The clean contigs were then structurally corrected using CRAQ v1.0.9-alpha [53] (default parameters) to break chimeric contigs, subsequently polished with Inspector v1.3.1 (--min_contig_length 100) [54]. Remaining chimeric contigs were broken using RagTag v2.1.0 (command: correct --remove-small --aligner nucmer --nucmer-params ‘--maxmatch -l 100 -c 500’ -v 45000) [55] with Darmor-bzh v10 serving as the reference genome. HiFi reads were employed in all correction steps. The corrected contigs were then scaffolded using RagTag v2.1.0 (-C -r -g 2 -m 9999999), also with Darmor-bzh v10 as reference. Assembly gaps were closed using TGSGapCloser v1.2.1 [56] without read correction (--ne). The continuity of the scaffolded assembly was assessed using gfastats v1.3.10. Chromosome lengths were calculated using bioawk v20110810. Gene space completeness was evaluated with BUSCO v5.8.3 (Embryophyta_odb12) [57]. Assembly correctness was assessed using the R-AQI and S-AQI metrics from CRAQ v1.0.9-alpha, as well as the quality value (QV) from Inspector v1.3.1. Structural correctness was further examined by aligning the assembly to Darmor-bzh v10 using D-Genies v1.5.0 (tool=minimap2; options=repeat:many) [58].

### Genome annotation and collinearity

Annotation was carried out using the RNAseq datasets generated in this study with Braker3 [59] (v3.0.8). Transposable elements were identified with EDTA [60] (v2.2.2) and annotated using repeatmasker [61] (v4.2.1). Gene ontology terms were transferred from Arabidopsis thaliana TAIR GO annotation to *B. napus* homologous genes, identified by matching protein sequences of *B. napus* and *A. thaliana* proteome database with BLASTp as previously described in [33, 62, 63]. Synteny analysis with MCScanX [64] was used to determine homoeolog pairs between the A and C subgenomes.

### ATAC data processing

ATACseq raw reads were trimmed with Cutadapt (v4.0) [65] and aligned to the newly generated reference (Express 617 v2) using Bowtie2 (v 2.4.5) [66] with the following settings: -k 10 -X 1000 --very-sensitive. Deeptools [67] was used to assess the correlation across biological replicates with multiBamSummary and plotCorrelation functions. Open chromatin regions or peaks were called with Genrich (https://github.com/harvardinformatics/Genrich) in ATAC mode (-j -y -a 200 -r -p 0.01). Peaks were called from both biological replicates directly with Genrich, which combined each replicates’ p-values by Fischer’s method. Peak annotation was carried out with ChIPseeker (v1.42) [68] and transcription factor binding sites were annotated with Meme FIMO (v 5.5.8) [69] with Jaspar database [70] and minimum adjusted pvalue (qvalue < 0.05).

### Super Enhancers identification

Super enhancers were defined as the top 2.5% non-near-promoter OCRs, as previously described in [24]. A custom bed file of all near promoters (500bp upstream of each TSS) was generated and used to remove all near-promoter OCRs with bedtools [71] intersect. Non-near-promoter OCRs length was used to sort them in decreasing size and the top 2.5% OCRs in each tissue were categorized as candidate super enhancers. Nearest (cognate) genes were identified with bedtools closest function (including distance in relation to the gene body with -D b). GO enrichment analysis was carried out with TopGO [72], using all OCR-cognate genes in the corresponding tissue/sample as background.

### WGBS data processing

WGBS raw reads were trimmed with trimgalore (v0.6.7) [73] with the following settings: --paired --clip_R1 8 --clip_R2 8 --three_prime_clip_R1 8 --three_prime_clip_R2 8. Methylation was called with Bismark v0.24.0 [74] against the Express 617 v3 reference, deduplicated with the deduplicate_bismark script and finally extracted in all three contexts with the bismark_methylation_extractor module. A minimum coverage threshold of five reads was applied for each cytosine for downstream analysis. Methylation levels across gene bodies, OCRs, distal OCRs were assessed with Deeptools computeMatrix and plotProfile.

### RNAseq data processing and ATAC-seq correlation

Raw RNAseq reads were trimmed with Cutadapt (-a CTGTCTCTTATACACATCT) and aligned with Hisat2 v2.2.1 [75] to the newly generated reference genome. Bam coverage was assessed with Deeptools (--normalizeUsing RPGC) for both RNAseq and ATACseq after bigwig conversion (bamCoverage) and averaging across replicates (bigwigAverage). Spearman correlation was calculated between the two data types by generating multibigwigsmmaries npz followed by plotCorrelation. Stringtie (v3.0.1) [76] was used to quantify transcripts abundances and summarize it at the gene level. Tissue specificity metrics (TAU) were calculated with tspex (v0.6.3) [77].

### SV calling and gene expression estimation in 57 additional *B. napus* genotypes

Following our approach from [33], long-read datasets (*n*: 57) were aligned using minimap2 v2.24-r1122 [78], followed by SV calling and genotyping with cuteSV v1.0.13 [79]. Variants were further filtered to retain homozygous insertions and deletions with genotype missing call rate: < 5%, MAF: > 5% and remove variants > 20 kb. Kallisto v0.44.0 [80] was used for expression quantification from the corresponding 57 genotypes using transcripts extracted from the Express 617 v3 assembly. TPM values were extracted directly from Kallisto outputs, and the combined expression matrix was transformed using inverse normal transformation.

### ML gene expression prediction integration

Gene expression was predicted with the *n*emo model based on the sequences of promoters and terminator regions annotated on the Express 617 v1 reference and compared to the observed expression across 8 tissues/ 36 RNA-seq samples [32]. Gene IDs were translated between references to match v1-based ML expression predictions to our v3-based SE-cognate genes. We set a minimum 99% of v3-based annotation overlapping minimum 50% v1-based annotation, to account for the lack of UTRs annotation in the v3 annotation approach.

### Weighted gene co-expression network analysis

Weighted gene co-expression network analysis (WGCNA) was performed with pyWGCNA (v2.0.0) with data from [27, 32] in addition to our newly generated RNAseq datasets. Genes were prefiltered to remove ones with low to no expression across all samples (<1 TPM) and the TPM matrix was used as input file for pyWGCNA preprocess() and findModule() functions.

## Acknowledgements

We thank the BRAVO project for granting us early access to their expression data for co-expression analysis.

## References

1. Schmitz RJ, Schultz MD, Urich MA, Nery JR, Pelizzola M, Libiger O, et al. Patterns of population epigenomic diversity. Nature. 2013;495:193–8. doi:10.1038/nature11968.

2. Crisp PA, Marand AP, Noshay JM, Zhou P, Lu Z, Schmitz RJ, Springer NM. Stable unmethylated DNA demarcates expressed genes and their cis-regulatory space in plant genomes. Proceedings of the National Academy of Sciences. 2020;117:23991–4000. doi:10.1073/pnas.2010250117.

3. Zhang H, Lang Z, Zhu J-K. Dynamics and function of DNA methylation in plants. Nat Rev Mol Cell Biol. 2018;19:489–506. doi:10.1038/s41580-018-0016-z.

4. Dowen RH, Pelizzola M, Schmitz RJ, Lister R, Dowen JM, Nery JR, et al. Widespread dynamic DNA methylation in response to biotic stress. Proc Natl Acad Sci U S A. 2012;109:E2183–91. doi:10.1073/pnas.1209329109.

5. Han Q, Bartels A, Cheng X, Meyer A, An Y-QC, Hsieh T-F, Xiao W. Epigenetics Regulates Reproductive Development in Plants. Plants. 2019;8:564. doi:10.3390/plants8120564.

6. Jason D. Buenrostro, Beijing Wu, Howard Y. Chang, William J. Greenleaf. ATAC-seq: A Method for Assaying Chromatin Accessibility Genome-Wide. Current Protocols in Molecular Biology. 2015;109:21.29.1-21.29.9. doi:10.1002/0471142727.mb2129s109.

7. Zhu T, Xia C, Yu R, Zhou X, Xu X, Wang L, et al. Comprehensive mapping and modelling of the rice regulome landscape unveils the regulatory architecture underlying complex traits. Nat Commun. 2024;15:6562. doi:10.1038/s41467-024-50787-y.

8. Feng D, Liang Z, Wang Y, Yao J, Yuan Z, Hu G, et al. Chromatin accessibility illuminates single-cell regulatory dynamics of rice root tips. BMC Biol. 2022;20:274. doi:10.1186/s12915-022-01473-2.

9. Liu Y, Gao X, Liu H, Yang X, Liu X, Xu F, et al. Constraint of accessible chromatins maps regulatory loci involved in maize speciation and domestication. Nat Commun. 2025;16:2477. doi:10.1038/s41467-025-57932-1.

10. Zhang W, Wu Y, Schnable JC, Zeng Z, Freeling M, Crawford GE, Jiang J. High-resolution mapping of open chromatin in the rice genome. Genome Res. 2012;22:151–62. doi:10.1101/gr.131342.111.

11. Wang M, Li Z, Zhang Y, Zhang Y, Xie Y, Ye L, et al. An atlas of wheat epigenetic regulatory elements reveals subgenome divergence in the regulation of development and stress responses. Plant Cell. 2021;33:865–81. doi:10.1093/plcell/koab028.

12. Yang H, Yu G, Lv Z, Li T, Wang X, Fu Y, et al. Epigenome profiling reveals distinctive regulatory features and cis-regulatory elements in pepper. Genome Biol. 2025;26:121. doi:10.1186/s13059-025-03595-6.

13. Lee H, Kang DH, Seo PJ. Enhancer Elements in Plants: Genomic Context and Biological Functions. J. Plant Biol. 2025;68:93–105. doi:10.1007/s12374-025-09460-0.

14. Yocca AE, Lu Z, Schmitz RJ, Freeling M, Edger PP. Evolution of Conserved Noncoding Sequences in Arabidopsis thaliana. Mol Biol Evol. 2021;38:2692–703. doi:10.1093/molbev/msab042.

15. Xu G, Lyu J, Li Q, Liu H, Wang D, Zhang M, et al. Evolutionary and functional genomics of DNA methylation in maize domestication and improvement. Nat Commun. 2020;11:5539. doi:10.1038/s41467-020-19333-4.

16. Eichten SR, Briskine R, Song J, Li Q, Swanson-Wagner R, Hermanson PJ, et al. Epigenetic and genetic influences on DNA methylation variation in maize populations. Plant Cell. 2013;25:2783–97. doi:10.1105/tpc.113.114793.

17. Zhang Z, Lin X, Yue J, Xu Y, Miao L, Tang W, et al. Reshaping epigenomic landscapes facilitated bread wheat speciation. Plant Physiol 2025. doi:10.1093/plphys/kiaf399.

18. Ding M, Chen ZJ. Epigenetic perspectives on the evolution and domestication of polyploid plant and crops. Curr Opin Plant Biol. 2018;42:37–48. doi:10.1016/j.pbi.2018.02.003.

19. Li H, Wang S, Chai S, Yang Z, Zhang Q, Xin H, et al. Graph-based pan-genome reveals structural and sequence variations related to agronomic traits and domestication in cucumber. Nat Commun. 2022;13:682. doi:10.1038/s41467-022-28362-0.

20. Bai F, Shu P, Deng H, Wu Y, Chen Y, Wu M, et al. A distal enhancer guides the negative selection of toxic glycoalkaloids during tomato domestication. Nat Commun. 2024;15:2894. doi:10.1038/s41467-024-47292-7.

21. Xie Y, Chen Y, Li Z, Zhu J, Liu M, Zhang Y, Dong Z. Enhancer transcription detected in the nascent transcriptomic landscape of bread wheat. Genome Biol. 2022;23:109. doi:10.1186/s13059-022-02675-1.

22. Fagny M, Kuijjer ML, Stam M, Joets J, Turc O, Rozière J, et al. Identification of Key Tissue-Specific, Biological Processes by Integrating Enhancer Information in Maize Gene Regulatory Networks. Front Genet. 2020;11:606285. doi:10.3389/fgene.2020.606285.

23. Zhu X, Chen A, Butler NM, Zeng Z, Xin H, Wang L, et al. Molecular dissection of an intronic enhancer governing cold-induced expression of the vacuolar invertase gene in potato. Plant Cell. 2024;36:1985–99. doi:10.1093/plcell/koae050.

24. Zhao H, Yang M, Bishop J, Teng Y, Cao Y, Beall BD, et al. Identification and functional validation of super-enhancers in Arabidopsis thaliana. Proceedings of the National Academy of Sciences. 2022;119:e2215328119. doi:10.1073/pnas.2215328119.

25. Hnisz D, Abraham BJ, Lee TI, Lau A, Saint-André V, Sigova AA, et al. Super-enhancers in the control of cell identity and disease. Cell. 2013;155:934–47. doi:10.1016/j.cell.2013.09.053.

26. Lee H, Chawla HS, Obermeier C, Dreyer F, Abbadi A, Snowdon R. Chromosome-Scale Assembly of Winter Oilseed Rape Brassica napus. Front. Plant Sci. 2020;11:514081. doi:10.3389/fpls.2020.00496.

27. Woolfenden H, Siles L, Vickers M, Steuernagel B, Morris RJ, Wells R, Kurup S. A dataset of tissue-specific gene expression dynamics during seed development in Brassica. Sci Data. 2025;12:800. doi:10.1038/s41597-025-05082-w.

28. Qing Zhang, Pengpeng Guan, Lun Zhao, Meng Ma, Liang Xie, Yue Li, et al. Asymmetric epigenome maps of subgenomes reveal imbalanced transcription and distinct evolutionary trends in Brassica napus. Molecular Plant. 2020;14:604–19. doi:10.1016/j.molp.2020.12.020.

29. Zhang Q, Guan P, Zhao L, Ma M, Xie L, Li Y, et al. Asymmetric epigenome maps of subgenomes reveal imbalanced transcription and distinct evolutionary trends in Brassica napus. Molecular Plant. 2021;14:604–19. doi:10.1016/j.molp.2020.12.020.

30. Mengdi Li, Weiqi Sun, Fan Wang, Xiaoming Wu, Jianbo Wang. Asymmetric epigenetic modification and homoeolog expression bias in the establishment and evolution of allopolyploid Brassica napus. New Phytologist. 2021;232:898–913. doi:10.1111/nph.17621.

31. Li Z, Li M, Wang J. Asymmetric subgenomic chromatin architecture impacts on gene expression in resynthesized and natural allopolyploid Brassica napus. Commun Biol. 2022;5:1–16. doi:10.1038/s42003-022-03729-7.

32. Kevin C. Rockenbach, Silvia F. Zanini, Richard J Morris, Rachel J Wells, Agnieszka A. Golicz. A deep learning model recapitulates position specific effects of plant regulatory sequences and suggests genes under complex regulation. bioRxiv. 2025:2025.08.30.673246. doi:10.1101/2025.08.30.673246.

33. Yildiz G, Zanini SF, Weber S, Kopalli V, Kox T, Abbadi A, et al. Graphical pangenomics-enabled characterization of structural variant impact on gene expression in Brassica napus. Theor Appl Genet. 2025;138:1–16. doi:10.1007/s00122-025-04867-2.

34. Grondin A, Rodrigues O, Verdoucq L, Merlot S, Leonhardt N, Maurel C. Aquaporins Contribute to ABA-Triggered Stomatal Closure through OST1-Mediated Phosphorylation. Plant Cell. 2015;27:1945–54. doi:10.1105/tpc.15.00421.

35. Tan Z, Peng Y, Xiong Y, Xiong F, Zhang Y, Guo N, et al. Comprehensive transcriptional variability analysis reveals gene networks regulating seed oil content of Brassica napus. Genome Biol. 2022;23:233. doi:10.1186/s13059-022-02801-z.

36. Zhang L, Liu L, Li H, He J, Chao H, Yan S, et al. 3D genome structural variations play important roles in regulating seed oil content of Brassica napus. Plant Comm. 2024;5:100666. doi:10.1016/j.xplc.2023.100666.

37. Zhang Y, Yang Z, He Y, Liu D, Liu Y, Liang C, et al. Structural variation reshapes population gene expression and trait variation in 2,105 Brassica napus accessions. Nat Genet. 2024;56:2538–50. doi:10.1038/s41588-024-01957-7.

38. Shan Tang, Hu Zhao, Shaoping Lu, Liangqian Yu, Guofang Zhang, Yuting Zhang, et al. Genome- and transcriptome-wide association studies provide insights into the genetic basis of natural variation of seed oil content in Brassica napus. Molecular Plant. 2020;14:470–87. doi:10.1016/j.molp.2020.12.003.

39. Zhang Y, Zhang H, Zhao H, Xia Y, Zheng X, Fan R, et al. Multi-omics analysis dissects the genetic architecture of seed coat content in Brassica napus. Genome Biol. 2022;23:1–22. doi:10.1186/s13059-022-02647-5.

40. Qu C, Zhao H, Fu F, Zhang K, Yuan J, Liu L, et al. Molecular Mapping and QTL for Expression Profiles of Flavonoid Genes in Brassica napus. Front. Plant Sci. 2016;7:210080. doi:10.3389/fpls.2016.01691.

41. He Y, Wu D, Wei D, Fu Y, Cui Y, Dong H, et al. GWAS, QTL mapping and gene expression analyses in Brassica napus reveal genetic control of branching morphogenesis. Sci Rep. 2017;7:1–9. doi:10.1038/s41598-017-15976-4.

42. Gajardo HA, Wittkop B, Soto-Cerda B, Higgins EE, Parkin IAP, Snowdon RJ, et al. Association mapping of seed quality traits in Brassica napus L. using GWAS and candidate QTL approaches. Mol Breeding. 2015;35:1–19. doi:10.1007/s11032-015-0340-3.

43. Cao J, Yu T, Xu B, Hu Z, Zhang X, Theurkauf WE, Weng Z. Epigenetic and chromosomal features drive transposon insertion in Drosophila melanogaster. Nucleic Acids Res. 2023;51:2066–86. doi:10.1093/nar/gkad054.

44. Wang M, Tu L, Lin M, Lin Z, Wang P, Yang Q, et al. Asymmetric subgenome selection and cis-regulatory divergence during cotton domestication. Nat Genet. 2017;49:579–87. doi:10.1038/ng.3807.

45. Yafang Xiao, Zengde Xi, Fei Wang, Jianbo Wang. Genomic asymmetric epigenetic modification of transposable elements is involved in gene expression regulation of allopolyploid Brassica napus. The Plant Journal. 2024;117:226–41. doi:10.1111/tpj.16491.

46. Dylan J. Ziegler, Deirdre Khan, Nadège Pulgar-Vidal, Isobel A. P. Parkin, Stephen J. Robinson, Mark F. Belmonte. Genomic asymmetry of the Brassica napus seed: epigenetic contributions of DNA methylation and small RNAs to subgenome bias. The Plant Journal. 2023;115:690–708. doi:10.1111/tpj.16254.

47. Wang F-X, Shang G-D, Wu L-Y, Mai Y-X, Gao J, Xu Z-G, Wang J-W. Protocol for assaying chromatin accessibility using ATAC-seq in plants. STAR Protoc. 2021;2:100289. doi:10.1016/j.xpro.2020.100289.

48. Lu Z, Hofmeister BT, Vollmers C, DuBois RM, Schmitz RJ. Combining ATAC-seq with nuclei sorting for discovery of cis-regulatory regions in plant genomes. Nucleic Acids Res. 2017;45:e41. doi:10.1093/nar/gkw1179.

49. Cheng H, Concepcion GT, Feng X, Zhang H, Li H. Haplotype-resolved de novo assembly using phased assembly graphs with hifiasm. Nat Methods. 2021;18:170–5. doi:10.1038/s41592-020-01056-5.

50. Wood DE, Lu J, Langmead B. Improved metagenomic analysis with Kraken 2. Genome Biol. 2019;20:257. doi:10.1186/s13059-019-1891-0.

51. Shen W, Ren H. TaxonKit: A practical and efficient NCBI taxonomy toolkit. Journal of Genetics and Genomics. 2021;48:844–50. doi:10.1016/j.jgg.2021.03.006.

52. The BLAST sequence analysis tool; 2013.

53. Li K, Xu P, Wang J, Yi X, Jiao Y. Identification of errors in draft genome assemblies at single-nucleotide resolution for quality assessment and improvement. Nat Commun. 2023;14:6556. doi:10.1038/s41467-023-42336-w.

54. Chen Y, Zhang Y, Wang AY, Gao M, Chong Z. Accurate long-read de novo assembly evaluation with Inspector. Genome Biol. 2021;22:312. doi:10.1186/s13059-021-02527-4.

55. Alonge M, Lebeigle L, Kirsche M, Jenike K, Ou S, Aganezov S, et al. Automated assembly scaffolding using RagTag elevates a new tomato system for high-throughput genome editing. Genome Biol. 2022;23:258. doi:10.1186/s13059-022-02823-7.

56. Xu M, Guo L, Gu S, Wang O, Zhang R, Peters BA, et al. TGS-GapCloser: A fast and accurate gap closer for large genomes with low coverage of error-prone long reads. Gigascience 2020. doi:10.1093/gigascience/giaa094.

57. Manni M, Berkeley MR, Seppey M, Simão FA, Zdobnov EM. BUSCO Update: Novel and Streamlined Workflows along with Broader and Deeper Phylogenetic Coverage for Scoring of Eukaryotic, Prokaryotic, and Viral Genomes. Mol Biol Evol. 2021;38:4647–54. doi:10.1093/molbev/msab199.

58. Cabanettes F, Klopp C. D-GENIES: dot plot large genomes in an interactive, efficient and simple way. PeerJ. 2018;6:e4958. doi:10.7717/peerj.4958.

59. Lars Gabriel, Tomáš Brůna, Katharina J. Hoff, Matthis Ebel, Alexandre Lomsadze, Mark Borodovsky, Mario Stanke. BRAKER3: Fully automated genome annotation using RNA-seq and protein evidence with GeneMark-ETP, AUGUSTUS, and TSEBRA. Genome Res. 2024;34:769–77. doi:10.1101/gr.278090.123.

60. Ou S, Su W, Liao Y, Chougule K, Agda JRA, Hellinga AJ, et al. Benchmarking transposable element annotation methods for creation of a streamlined, comprehensive pipeline. Genome Biol. 2019;20:1–18. doi:10.1186/s13059-019-1905-y.

61. Smit AF, Hubley R, Green P. RepeatMasker Open-4.0. 2013–2015 2015: Seattle, USA.

62. Golicz AA, Allu AD, Li W, Lohani N, Singh MB, Bhalla PL. A dynamic intron retention program regulates the expression of several hundred genes during pollen meiosis. Plant Reproduction. 2021;34:225–42. doi:10.1007/s00497-021-00411-6.

63. Sessa EB, Masalia RR, Arrigo N, Barker MS, Pelosi JA. GOgetter: A pipeline for summarizing and visualizing GO slim annotations for plant genetic data. Appl. Plant Sci. 2023;11:e11536. doi:10.1002/aps3.11536.

64. Wang Y, Tang H, Wang X, Sun Y, Joseph PV, Paterson AH. Detection of colinear blocks and synteny and evolutionary analyses based on utilization of MCScanX. Nat Protoc. 2024;19:2206–29. doi:10.1038/s41596-024-00968-2.

65. Marcel Martin. Cutadapt removes adapter sequences from high-throughput sequencing reads. EMBnet j. 2011;17:10–2. doi:10.14806/ej.17.1.200.

66. Langmead B, Salzberg SL. Fast gapped-read alignment with Bowtie 2. Nat Methods. 2012;9:357–9. doi:10.1038/nmeth.1923.

67. Ramírez F, Dündar F, Diehl S, Grüning BA, Manke T. deepTools: a flexible platform for exploring deep-sequencing data. Nucleic Acids Res. 2014;42:W187–W191. doi:10.1093/nar/gku365.

68. Yu G, Wang L-G, He Q-Y. ChIPseeker: an R/Bioconductor package for ChIP peak annotation, comparison and visualization. Bioinformatics. 2015;31:2382–3. doi:10.1093/bioinformatics/btv145.

69. Grant CE, Bailey TL, Noble WS. FIMO: scanning for occurrences of a given motif. Bioinformatics. 2011;27:1017–8. doi:10.1093/bioinformatics/btr064.

70. Sandelin A, Alkema W, Engström P, Wasserman WW, Lenhard B. JASPAR: an open-access database for eukaryotic transcription factor binding profiles. Nucleic Acids Res. 2004;32:D91–D94. doi:10.1093/nar/gkh012.

71. Quinlan AR, Hall IM. BEDTools: a flexible suite of utilities for comparing genomic features. Bioinformatics. 2010;26:841–2. doi:10.1093/bioinformatics/btq033.

72. Adrian Alexa JR. topGO: Bioconductor; 2017.

73. Krueger F. Trim Galore!: A wrapper around Cutadapt and FastQC to consistently apply adapter and quality trimming to FastQ files, with extra functionality for RRBS data. Babraham Institute. 2015.

74. Krueger F, Andrews SR. Bismark: a flexible aligner and methylation caller for Bisulfite-Seq applications. Bioinformatics. 2011;27:1571–2. doi:10.1093/bioinformatics/btr167.

75. Kim D, Paggi JM, Park C, Bennett C, Salzberg SL. Graph-based genome alignment and genotyping with HISAT2 and HISAT-genotype. Nat Biotechnol. 2019;37:907–15. doi:10.1038/s41587-019-0201-4.

76. Pertea M, Pertea GM, Antonescu CM, Chang T-C, Mendell JT, Salzberg SL. StringTie enables improved reconstruction of a transcriptome from RNA-seq reads. Nat Biotechnol. 2015;33:290–5. doi:10.1038/nbt.3122.

77. Antonio P. Camargo, Adrielle A. Vasconcelos, Mateus B. Fiamenghi, Gonçalo A. G. Pereira, Marcelo F. Carazzolle. tspex: a tissue-specificity calculator for gene expression data 2020. doi:10.21203/rs.3.rs-51998/v1.

78. Li H. Minimap2: pairwise alignment for nucleotide sequences. Bioinformatics. 2018;34:3094–100. doi:10.1093/bioinformatics/bty191.

79. Jiang T, Liu Y, Jiang Y, Li J, Gao Y, Cui Z, et al. Long-read-based human genomic structural variation detection with cuteSV. Genome Biol. 2020;21:189. doi:10.1186/s13059-020-02107-y.

80. Bray NL, Pimentel H, Melsted P, Pachter L. Near-optimal probabilistic RNA-seq quantification. Nat Biotechnol. 2016;34:525–7. doi:10.1038/nbt.3519.

