## Supplemental Material for "Integrative epigenomic analysis uncovers asymmetry of enhancer activity in *Brassica napus*"

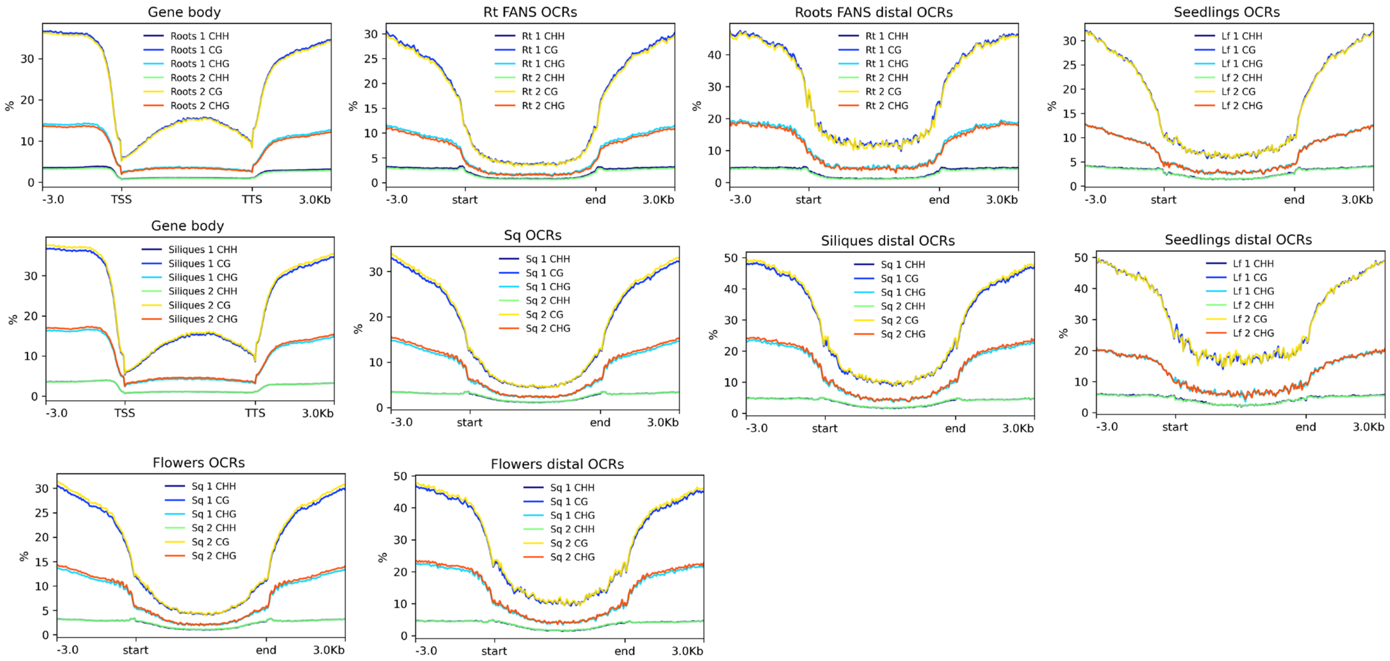


Supplementary figure 1. DNA methylation in all 3 contexts (CHH,CHG and CG) across genic regions , OCRs and distal OCRs in roots, siliques , seedlings and flowers

Supplementary Figure 2 Pearson correlation between ATACseq and  mRNAseq bam coverages across extended genes (gene body + promoter (3kb))

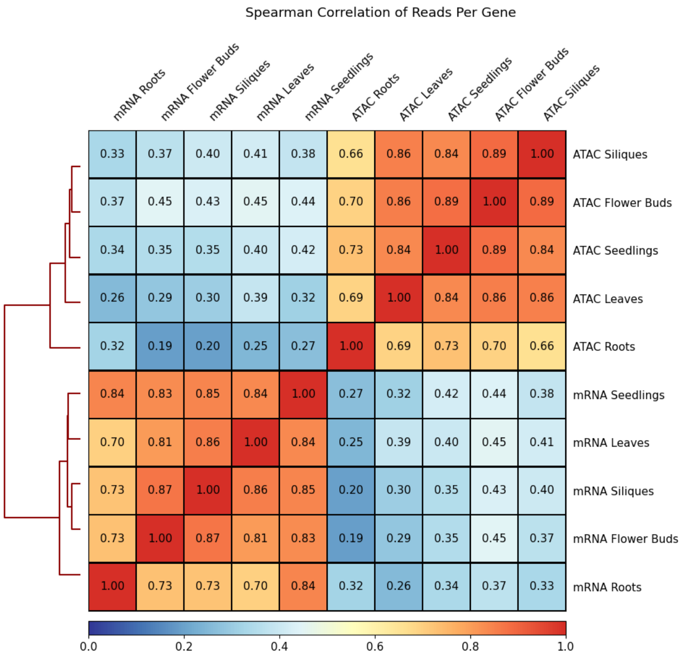

Supplementary Figure 3 top Gene Ontology terms enriched in SE-cognate genes in roots, seedlings and flowers.

| Roots SEs | | | |
|---|---|---|---|
| **GO** | **Term** | **# of Genes** | **P-value** |
| GO:0015986 | proton motive force-driven ATP synthesis | 4 | 3.20E-04 |
| GO:0006536 | glutamate metabolic process | 4 | 3.70E-04 |
| GO:0046777 | protein autophosphorylation | 8 | 8.40E-04 |
| GO:0019725 | cellular homeostasis | 8 | 2.14E-03 |
| GO:0006979 | response to oxidative stress | 13 | 3.71E-03 |
| GO:0050832 | defense response to fungus | 8 | 4.76E-03 |
| GO:0010118 | stomatal movement | 7 | 1.71E-02 |
| GO:0071456 | cellular response to hypoxia | 9 | 2.13E-02 |
| GO:0009968 | negative regulation of signal transducti... | 5 | 4.00E-02 |
| GO:0009723 | response to ethylene | 7 | 4.70E-02 |

| Seedlings SEs | | | |
|---|---|---|---|
| **GO** | **Term** | **# of Genes** | **P-value** |
| GO:0042547 | cell wall modification involved in multi... | 7 | 3.00E-08 |
| GO:0009606 | tropism | 10 | 2.00E-06 |
| GO:0009827 | plant-type cell wall modification | 8 | 3.60E-06 |
| GO:0009960 | endosperm development | 5 | 2.50E-05 |
| GO:0009629 | response to gravity | 8 | 2.70E-05 |
| GO:0010143 | cutin biosynthetic process | 4 | 1.50E-04 |
| GO:0018107 | peptidyl-threonine phosphorylation | 4 | 1.50E-04 |
| GO:0046777 | protein autophosphorylation | 7 | 6.40E-04 |
| GO:0015986 | proton motive force-driven ATP synthesis | 4 | 1.44E-03 |
| GO:0010214 | seed coat development | 5 | 1.81E-03 |

| Flowers SEs | | | |
|---|---|---|---|
| **GO** | **Term** | **# of Genes** | **P-value** |
| GO:0010214 | seed coat development | 14 | 7.90E-10 |
| GO:2000033 | regulation of seed dormancy process | 7 | 9.30E-08 |
| GO:0051093 | negative regulation of developmental pro... | 20 | 3.60E-07 |
| GO:0048833 | specification of floral organ number | 5 | 4.30E-07 |
| GO:0009825 | multidimensional cell growth | 10 | 5.10E-07 |
| GO:0009827 | plant-type cell wall modification | 11 | 1.80E-06 |
| GO:0015936 | coenzyme A metabolic process | 6 | 2.80E-06 |
| GO:0051241 | negative regulation of multicellular org... | 16 | 5.90E-06 |
| GO:0009968 | negative regulation of signal transducti... | 15 | 8.00E-06 |
| GO:0042447 | hormone catabolic process | 4 | 1.20E-05 |

Supplementary Figure 4 median gene connectivity values across 29 gene modules based on genes' proximity to open chromatin regions, super-enhancers or neither. Significance test: Mann-Whitney U test.

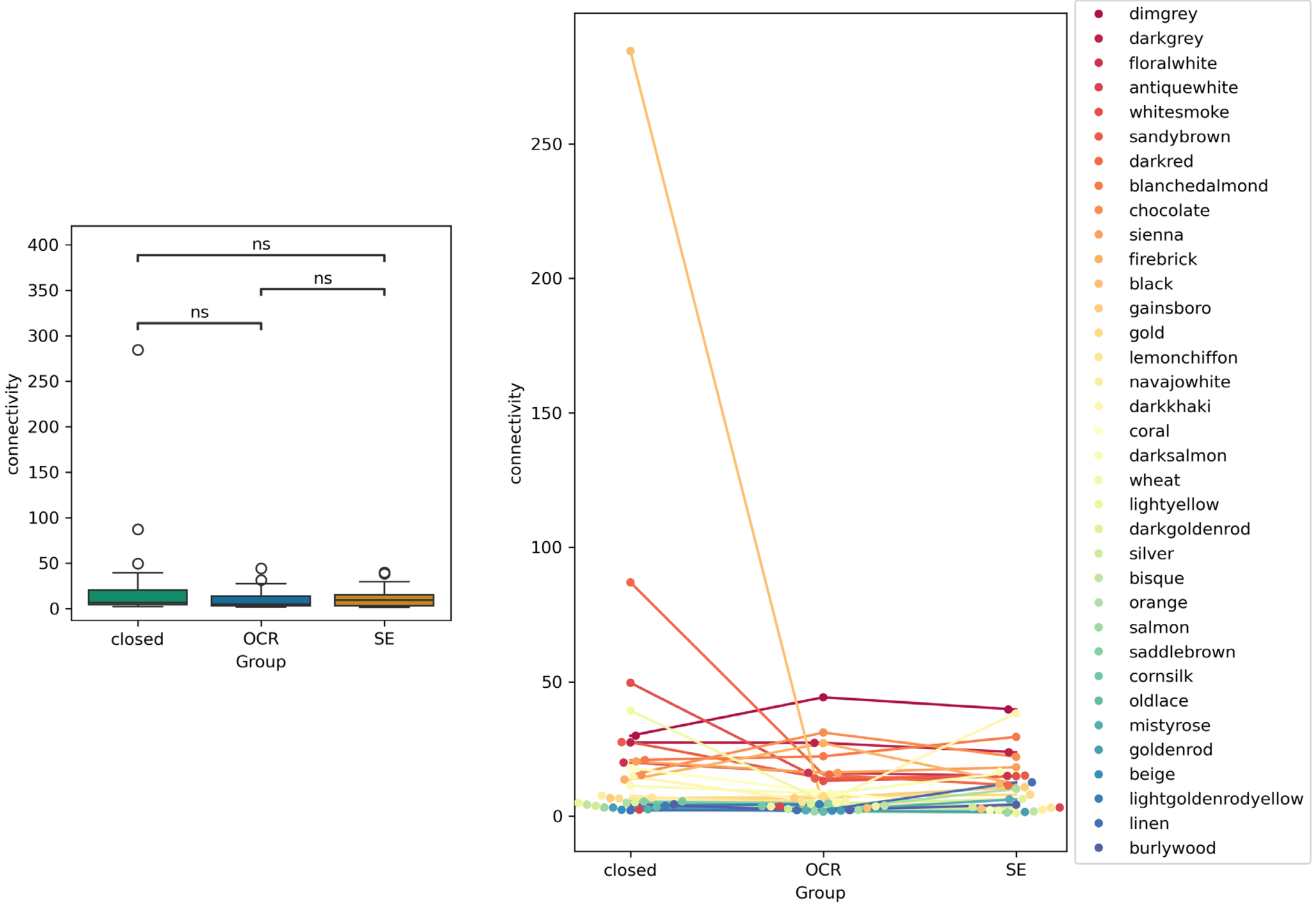

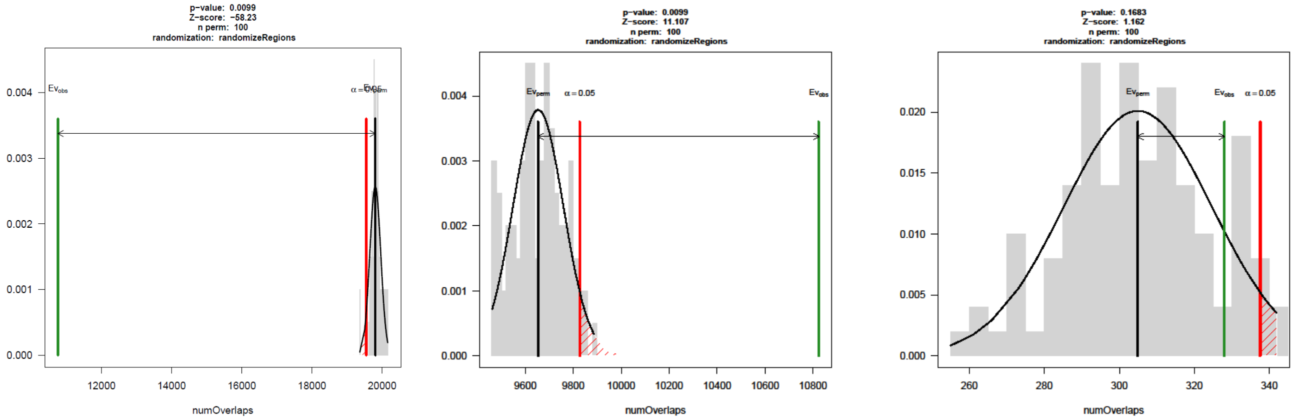


Supplementary Figure 5 regioneR permutation tests for assessing the enrichment or depletion of SVs across a) genes, b) open chromatin regions and c) super-enhancers.

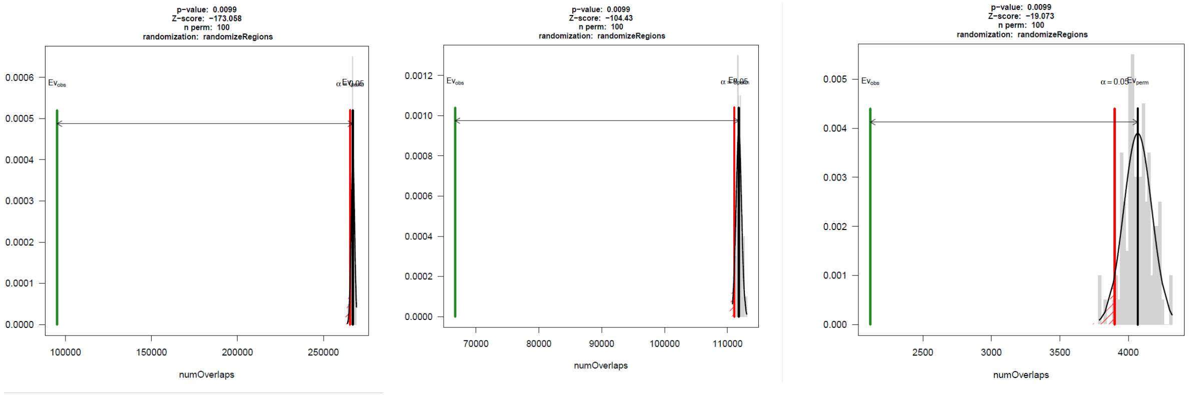


Supplementary Figure 6 regioneR permutation tests for assessing the enrichment or depletion of transposable elements (TEs) across a) genes, b) open chromatin regions and c) super-enhancers.
